## Supplementary Data for "Atomic resolution structures of the methane-activating enzyme in anaerobic methanotrophy reveal extensive post-translational modifications"

### Supplementary material.

**This file contains the Supplementary Figures S1-18 and references.**

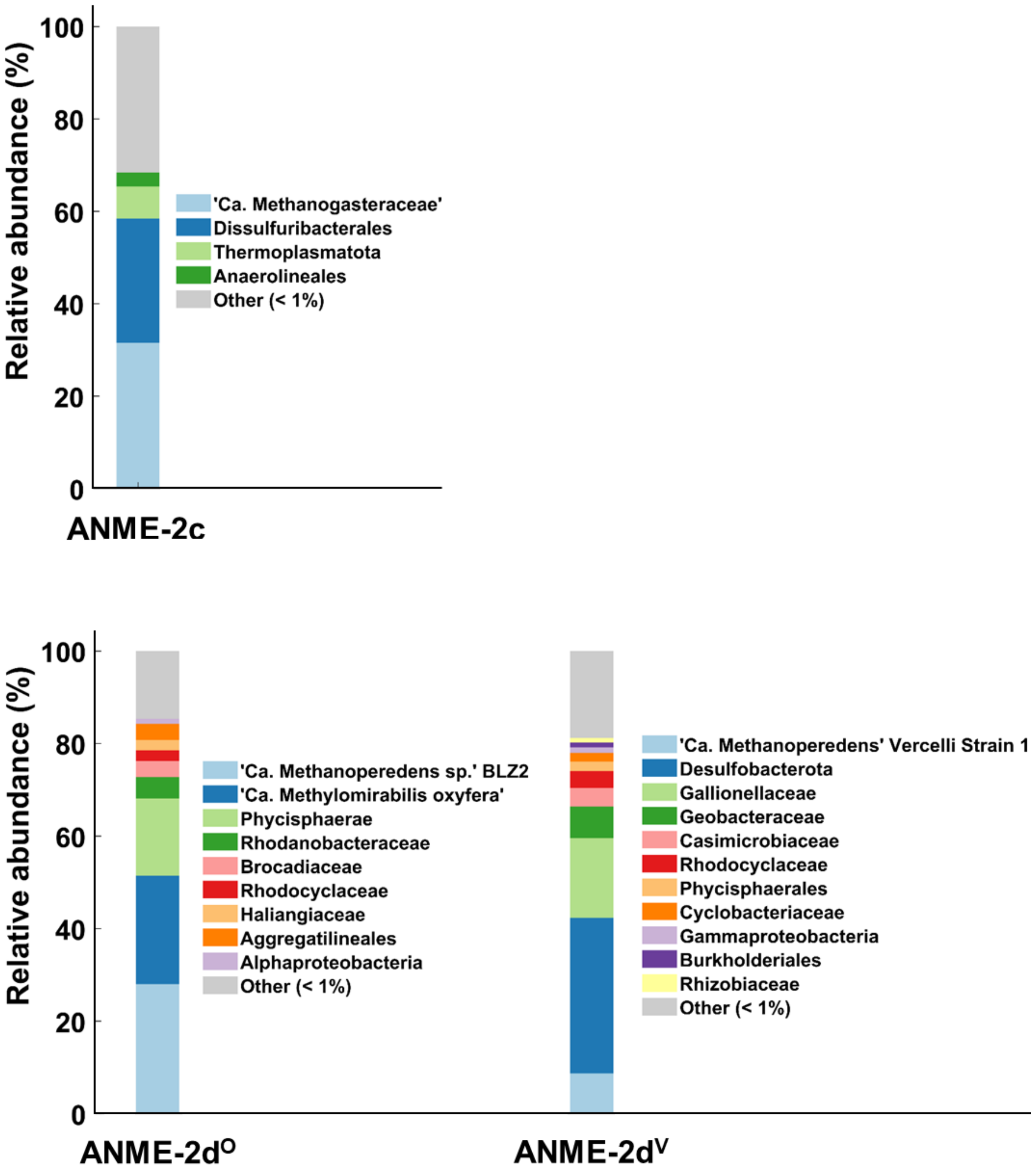


**Supplementary Figure S1.** Metagenomic read-based relative abundances of ANME enrichments. SingleM was used for taxonomical read-classification on three ANME enrichments using the GTDB-Tk database version 2.4.0 with raw reads as input(1).

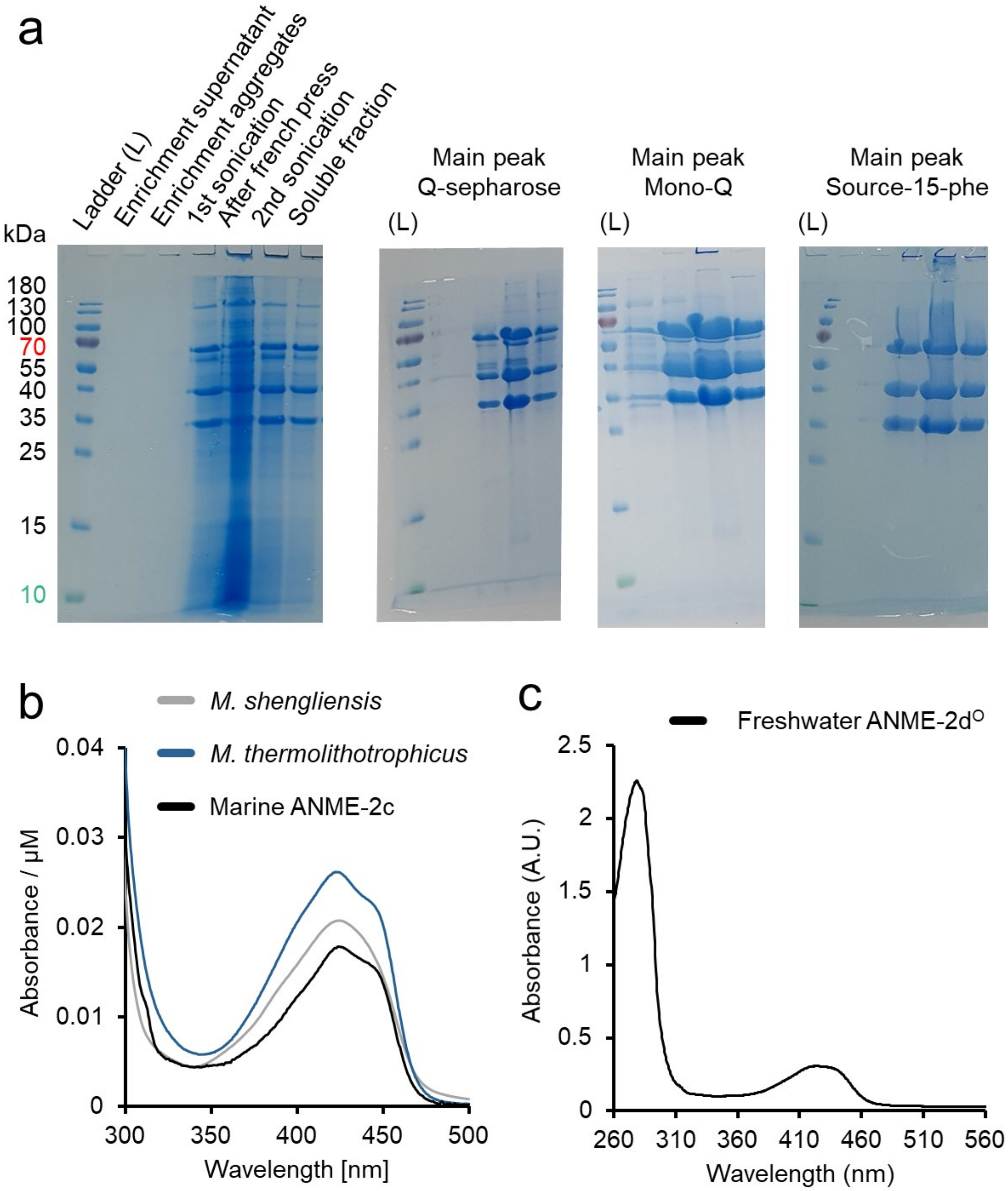


**Supplementary Figure S2. Purification of ANME-2c MCR and UV/Visible spectroscopy of studied samples**. (**a**) SDS-PAGE of different purification steps. L stands for Ladder. (**b**) UV-visible spectrum of ANME-2c MCR compared to methanogenic homologs from *Methermicoccus shengliensis*(2) and *Methanothermococcus thermolithotrophicus*(3). The measured signal is in absorbance (AU) per µM of measured MCR. (**c**) UV-visible spectrum of ANME-2d$^{O}$ MCR.

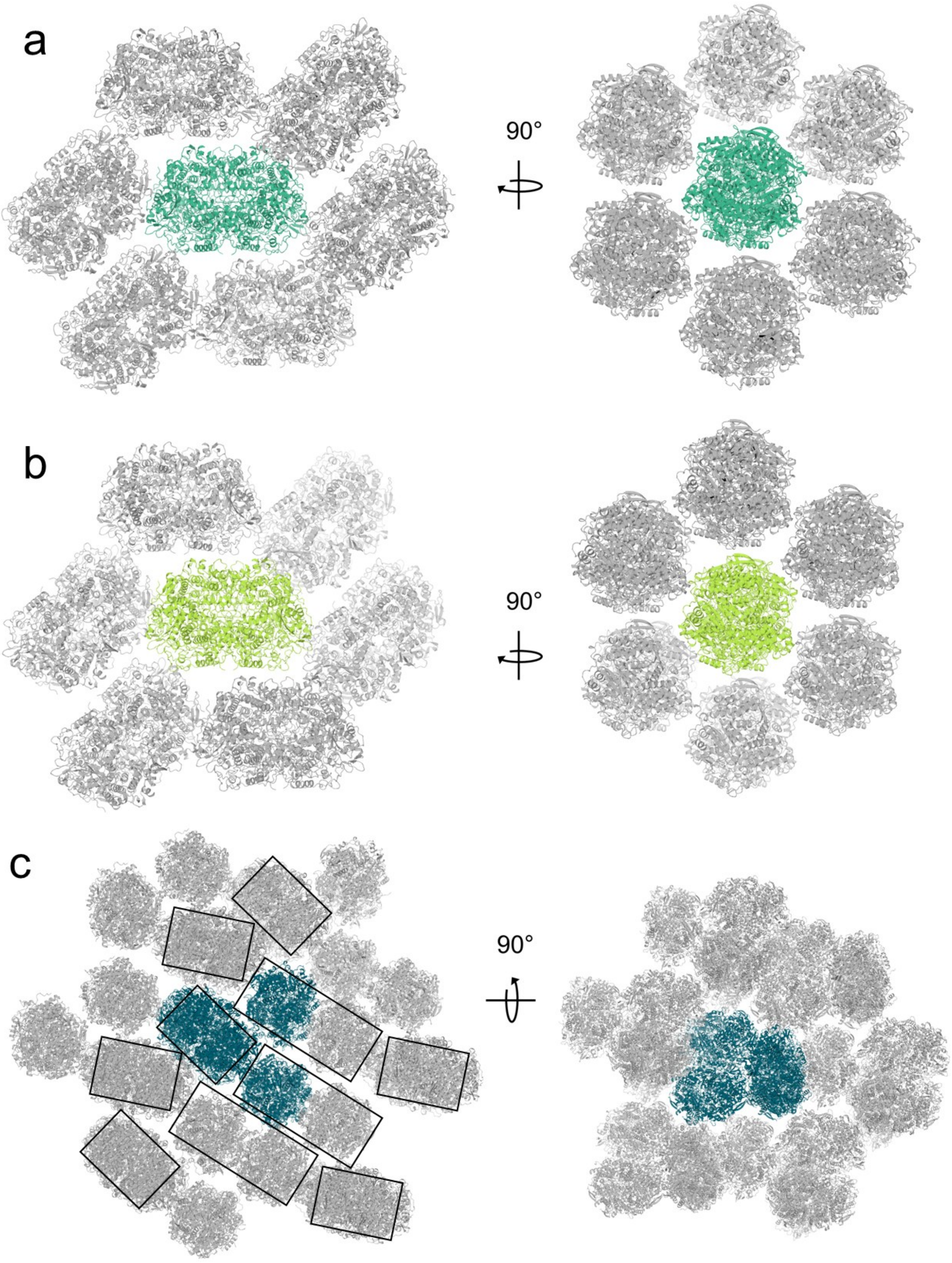


**Supplementary Figure S3. Presentation of ANME-2 MCR crystalline packing.** Packing of MCR from ANME-2d$^{O}$ (**a**), ANME-2d$^{V}$ (**b**), and ANME-2c (**c**). MCR units are shown in cartoons. The asymmetric unit content is coloured, while the symmetry mates are grey. Black boxes indicate observable dimeric units for ANME-2c MCR, which might come from a crystallisation artefact.

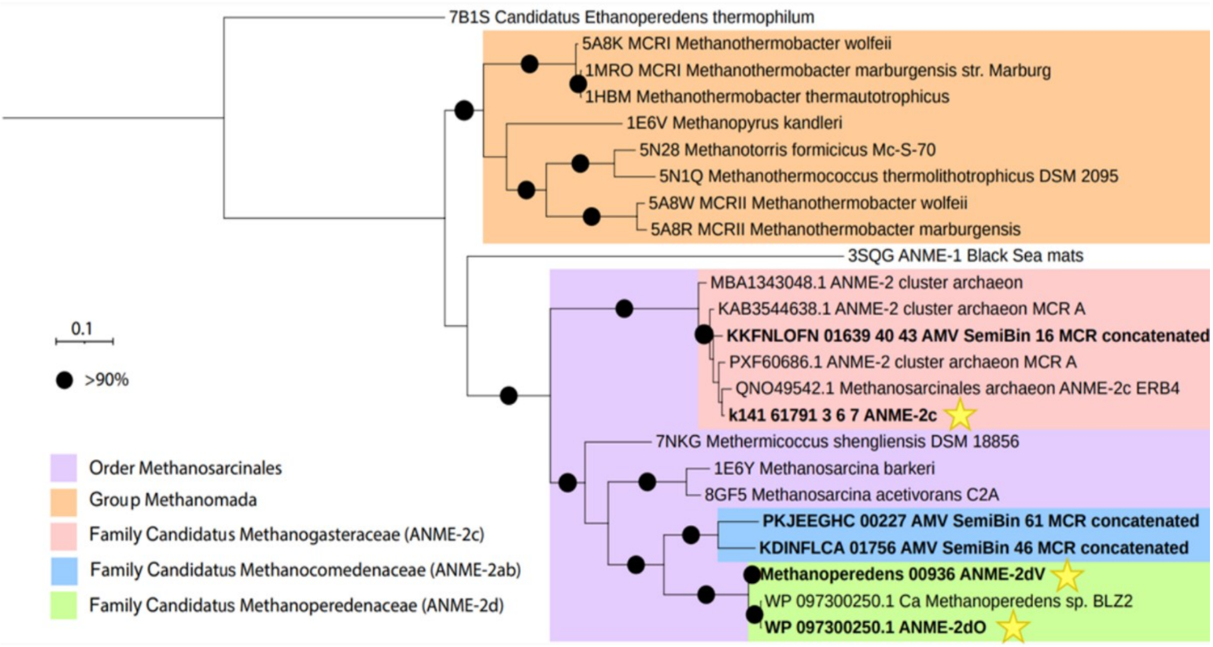


**Supplementary Figure S4. Phylogeny tree of the structurally characterised MCRs and close sequence homologues from the marine ANME-2 MCR.** The tree represents the concatenation of the α, β, and γ MCR subunits. The models studied in this work are highlighted with yellow stars.

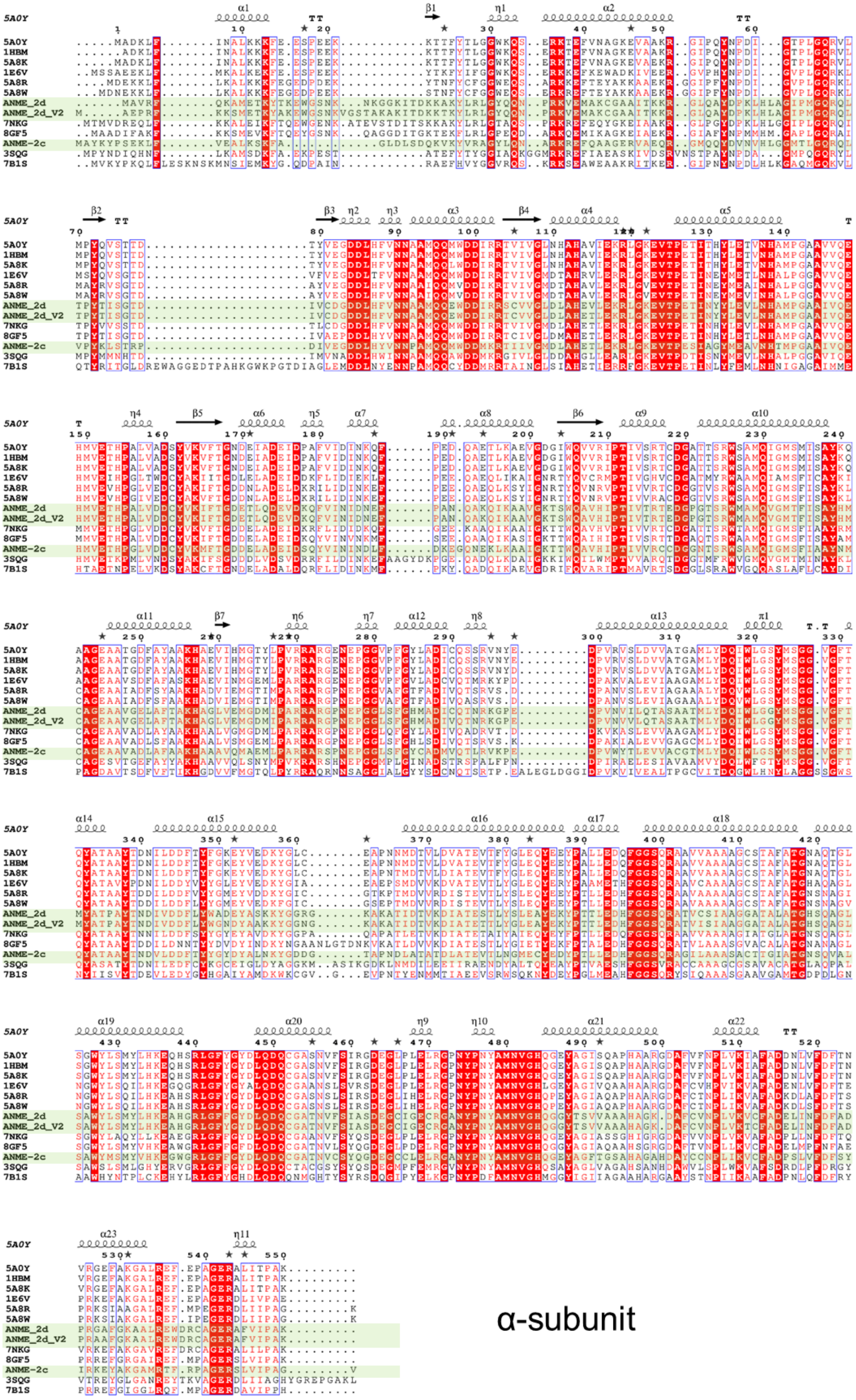

α-subunit

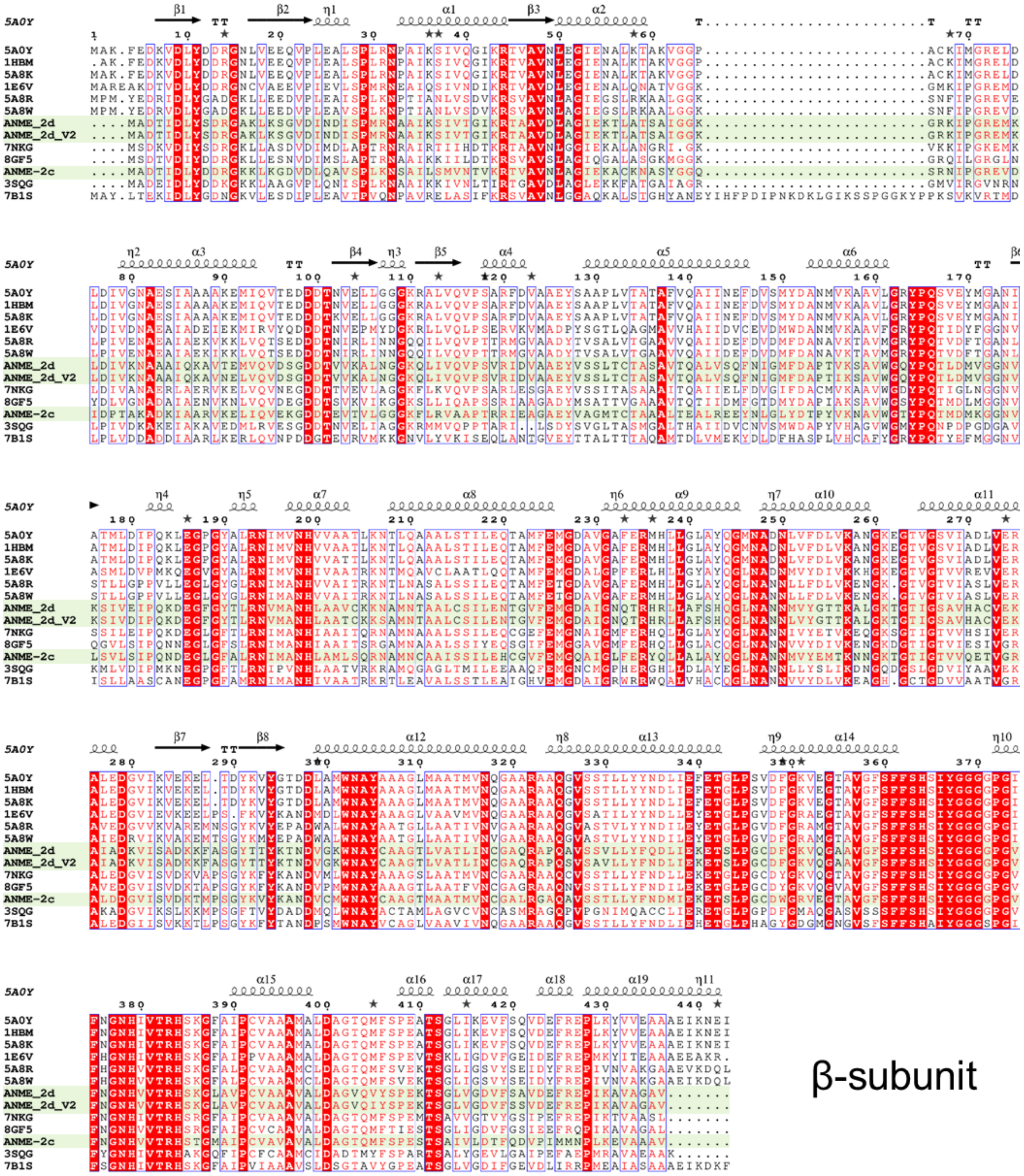
5A0Y
1HBM
5A8K
1E6V
5A8R
5A8W
ANME_2d
ANME_2d_V2
7NKG
8GF5
ANME-2c
3SQG
7B1S
β-subunit

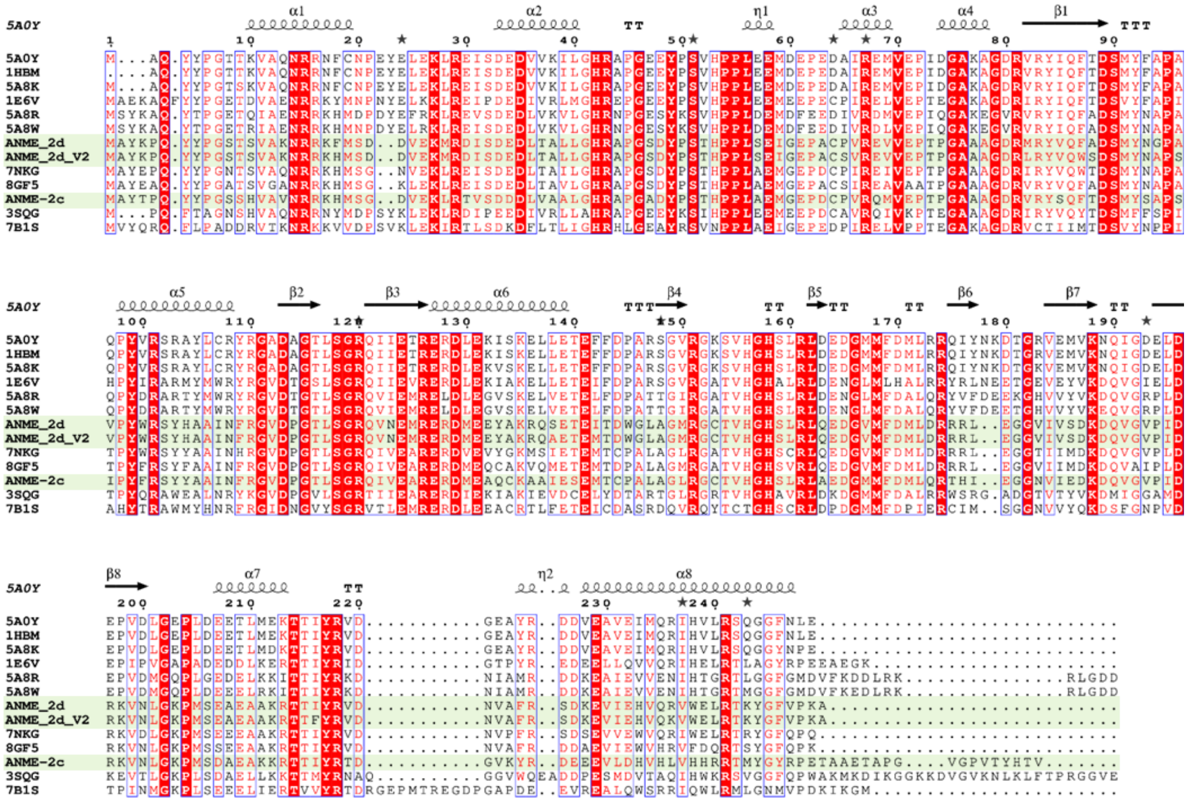


γ-subunit

**Supplementary Figure S5. Sequence alignment of the structurally characterised MCR homologs.** ESPript alignment(4) of marine ANME-2c, freshwater ANME-2d$^{O}$ (ANME_2d), freshwater ANME-2d$^{V}$ (ANME_2d_V2) and structurally characterised homologs from *Methanothermobacter marburgensis* isoform I (PDB code: 5A0Y), *Methanothermobacter thermautotrophicus* (1HBM), *Methanothermobacter wolfeii* isoform I (5A8K), *Methanopyrus kandleri* (1E6V), *Methanothermobacter marburgensis* isoform II (5A8R), *Methanothermobacter wolfeii* isoform II (5A8W), *Methermicoccus shengliensis* (7NKG), *Methanosarcina acetivorans* (8GF5), ANME-1 from Black Sea mats (3SQG) and '*Candidatus* Ethanoperedens thermophilum' (7B1S). ANME-2 MCR sequences are highlighted in green.

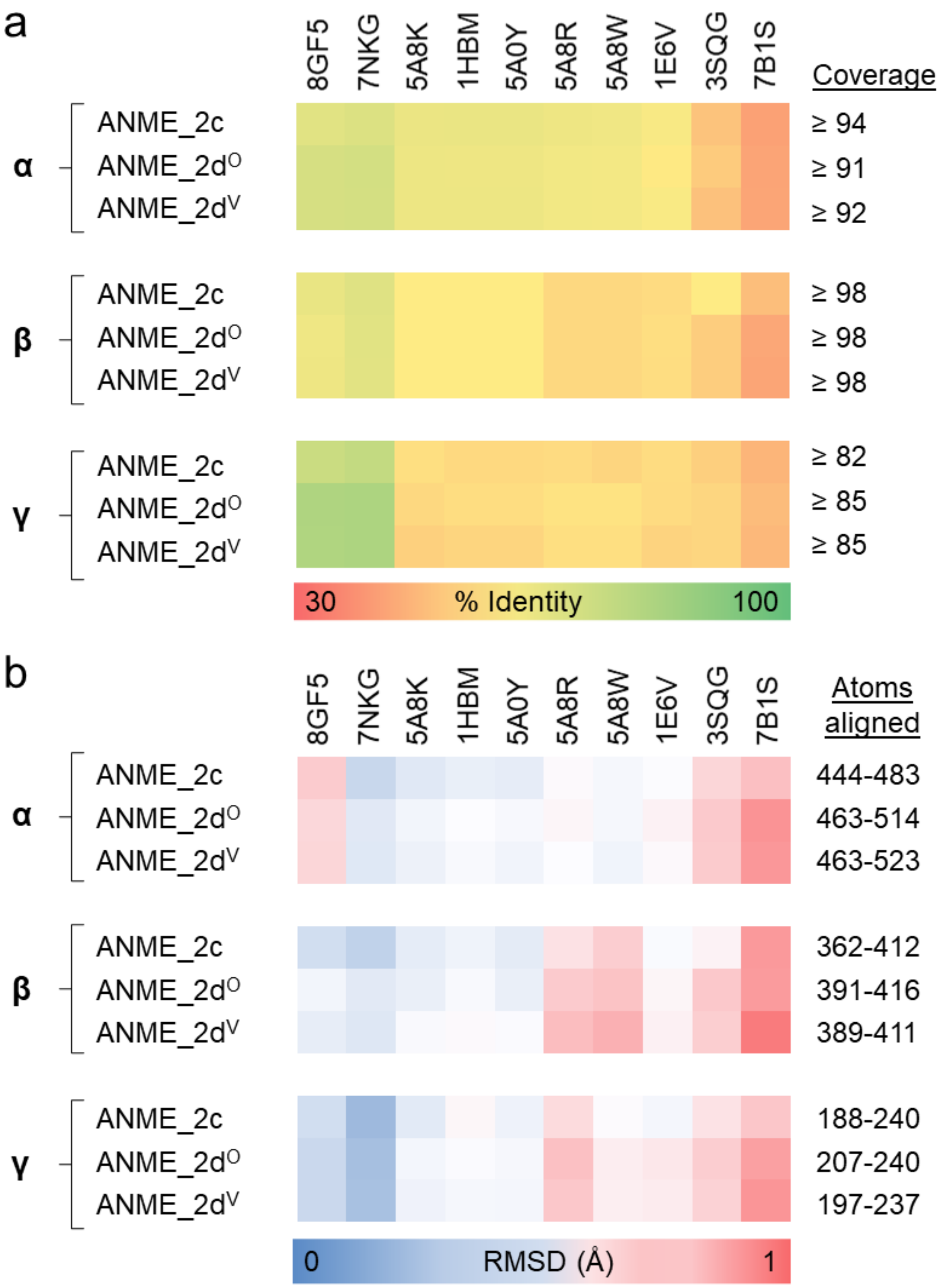


**Supplementary Figure S6. Sequence and structural similarity of characterised MCR homologs.** (**a**) Sequence similarity of the α-, β- and γ-subunits of ANME-2 MCRs to structural homologs. The percentage of identity is displayed as a gradient from red to green, ranging from 30% to 100% sequence identity. The coverage for all samples is summarised on the right. (**b**) Structural similarity of the α-, β- and γ-subunits of ANME-2 MCRs to structural homologs. The RMSD is displayed as a gradient from blue to red, ranging from 0 to 1 Å. The aligned Cα atoms for all samples are summarised on the right. Listed homologs are *M. marburgensis* isoform I (PDB code: 5A0Y), *M. thermautotrophicus* (1HBM), *M. wolfeii* isoform I (5A8K), *M. kandleri* (1E6V), *M. marburgensis* isoform II (5A8R), *M. wolfeii* isoform II (5A8W), *M. shengliensis* (7NKG), *M. acetivorans* (8GF5), ANME-1 (3SQG), and '*Ca*. E. thermophilum' (7B1S).

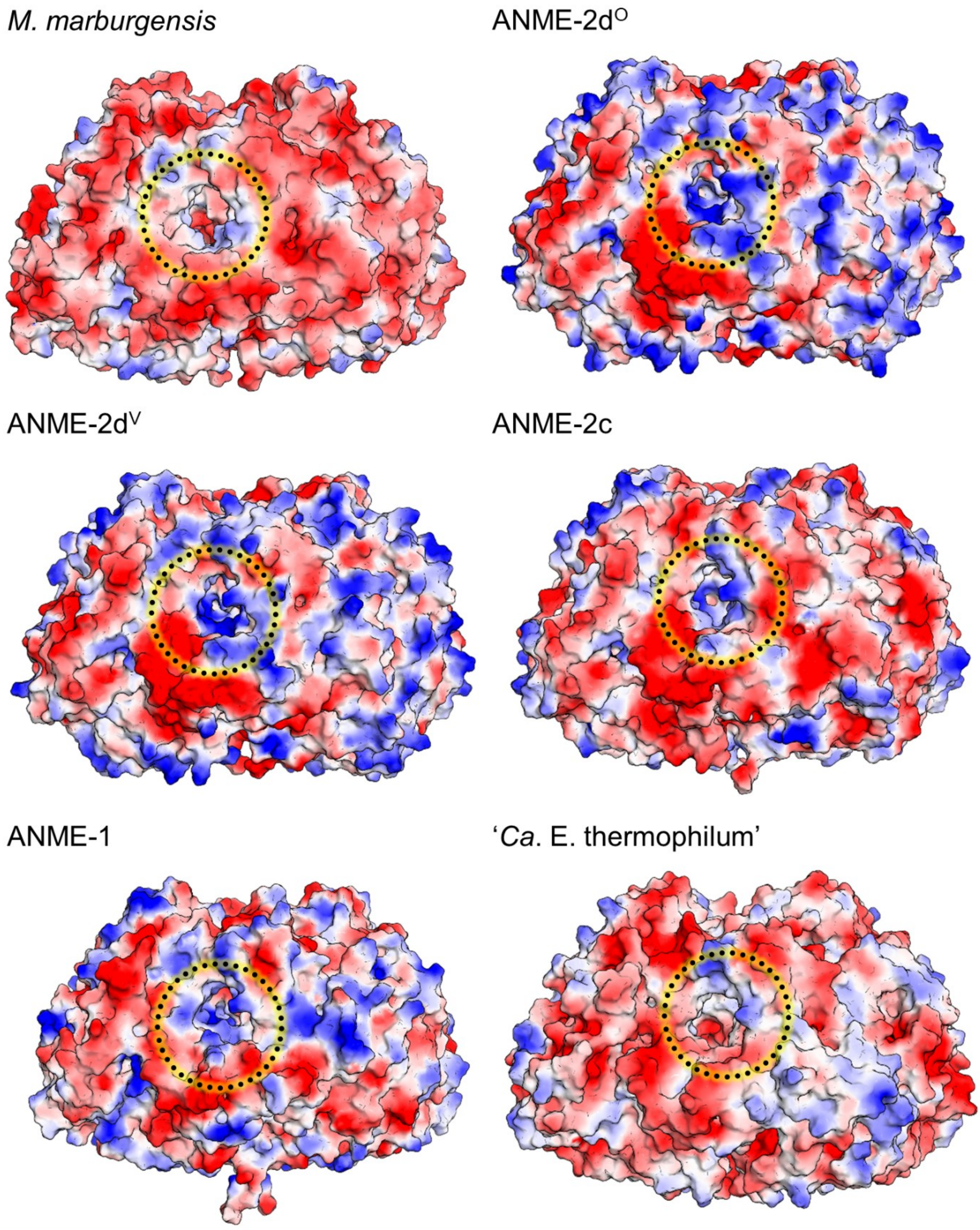


**Supplementary Figure S7. Electrostatic potential on the surface of MCRs.** The vacuum electrostatic potential of MCR is shown as a gradient from red to blue for a more negative to a positive potential, respectively. Dashed circles with a yellow glow highlight the active site entrance. PDB codes for the models are the following: *M. marburgensis* isoform I (PDB code: 5A0Y), *M. shengliensis* (7NKG), ANME-1 (3SQG), and '*Ca*. E. thermophilum' (7B1S).

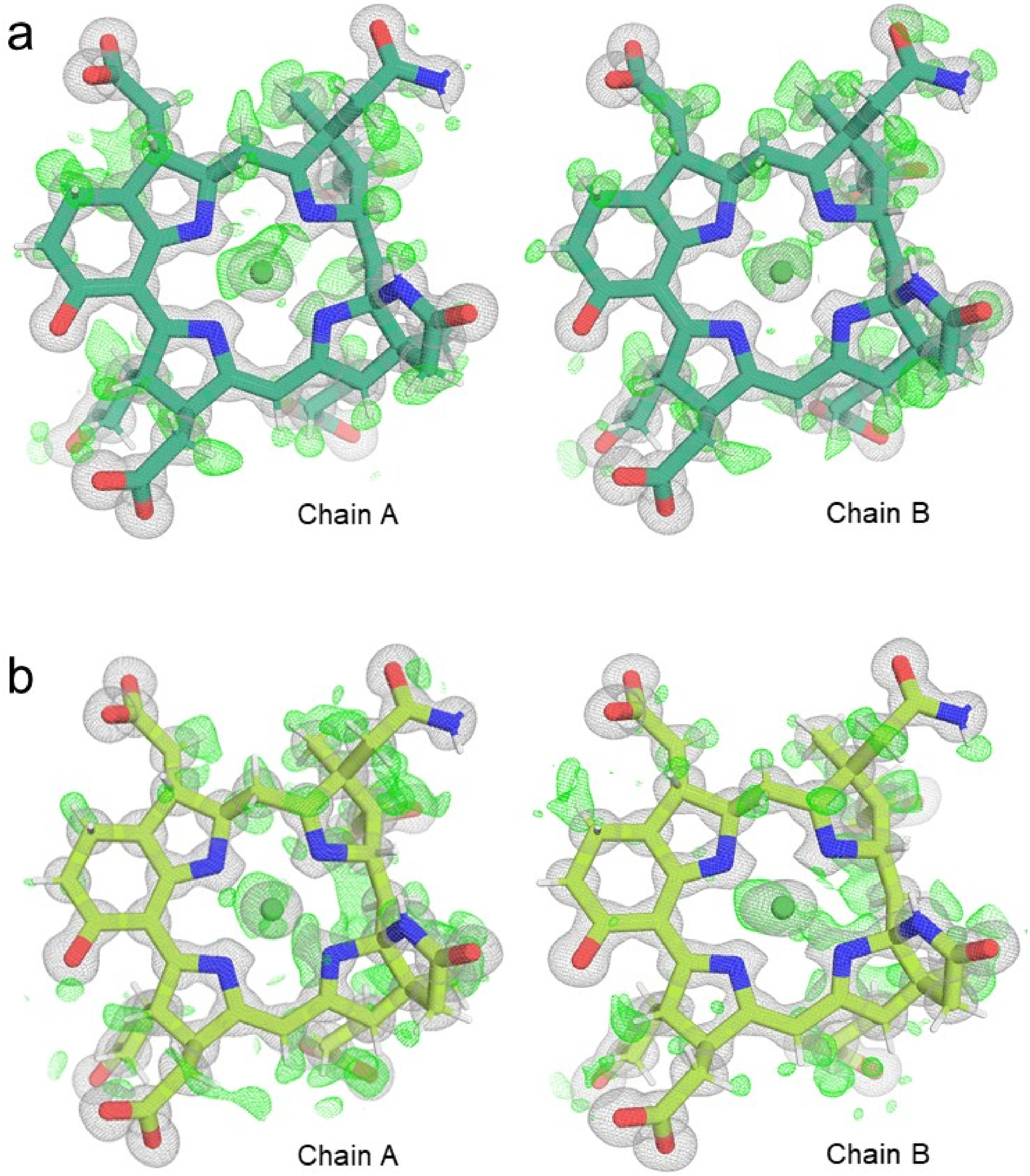


**Supplementary Figure S8. Electron density map of $F_{430}$.** The electron density is shown for (**a**) ANME-2d$^{O}$ MCR and (**b**) ANME-2d$^{V}$ MCR as grey mesh for the $2F_o{-}F_c$ map at 1 σ and green mesh for the $F_o{-}F_c$ map at 2.8 σ. $F_{430}$ is shown as sticks. Atoms are coloured in red for O, blue for N, and white for riding H. The map was generated with non-hydrogenated $F_{430}$s.

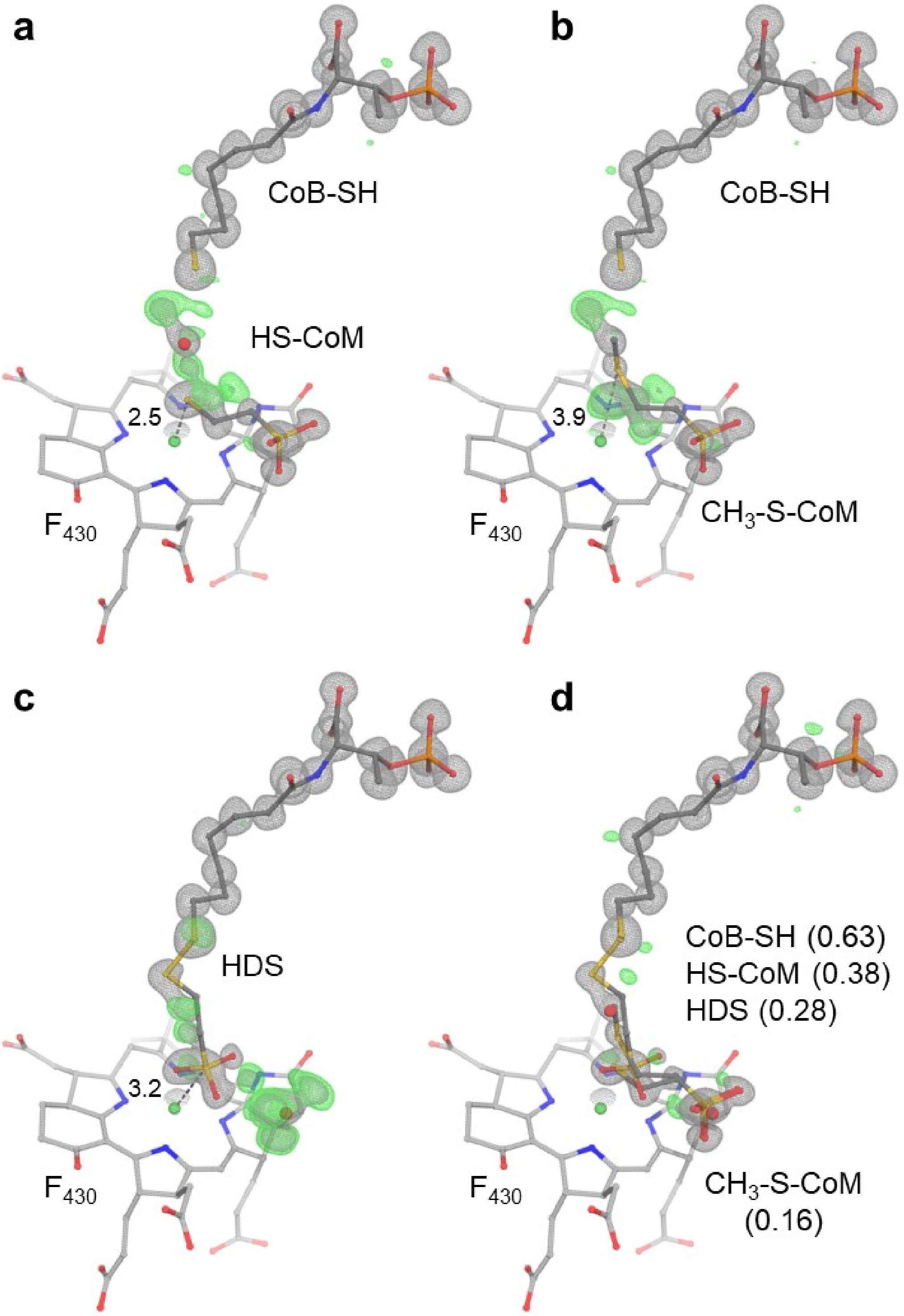


**Supplementary Figure S9. Ligand modelling in ANME-2d$^{V}$ MCR active site.** Ligand mixtures were modelled as follows: (**a**) Coenzyme B and coenzyme M, (**b**) coenzyme B and methyl-coenzyme M, (**c**) heterodisulfide of coenzyme M and coenzyme B and (**d**) all four ligands. Models were refined with ligands at 1.0 occupancy except for (**d**), where partial occupancies are indicated in brackets. Ligands are shown as sticks with atoms coloured in red for O, blue for N, yellow for S, green for Ni and orange for P. The $2F_o{-}F_c$ map is contoured at 1 σ and the $F_o{-}F_c$ map is contoured at 4 σ and shown as grey and green mesh, respectively. The distance (Å) between the S and Ni atom is shown as black dashes.

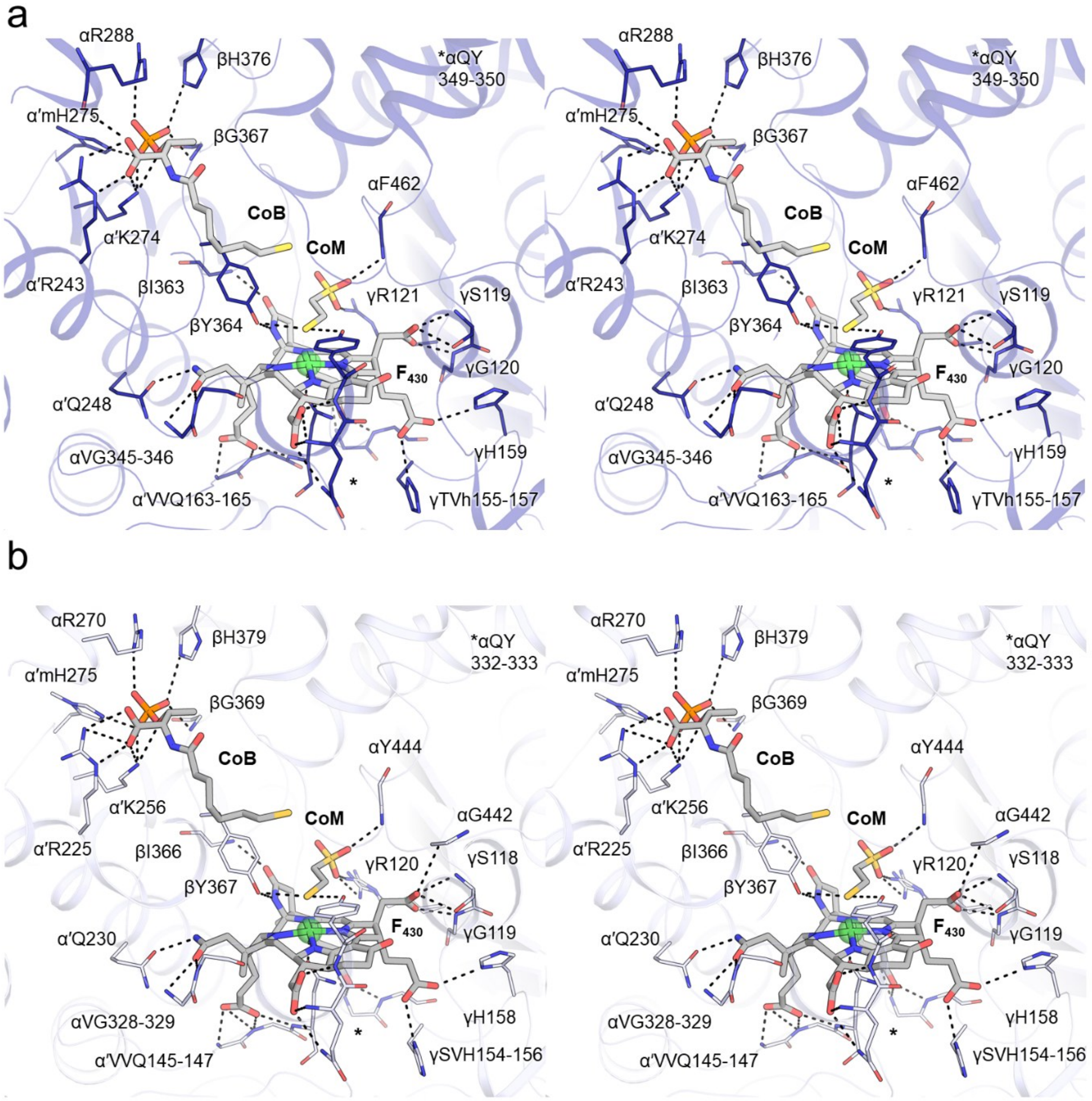


**Supplementary Figure S10.** Continued on the next page.

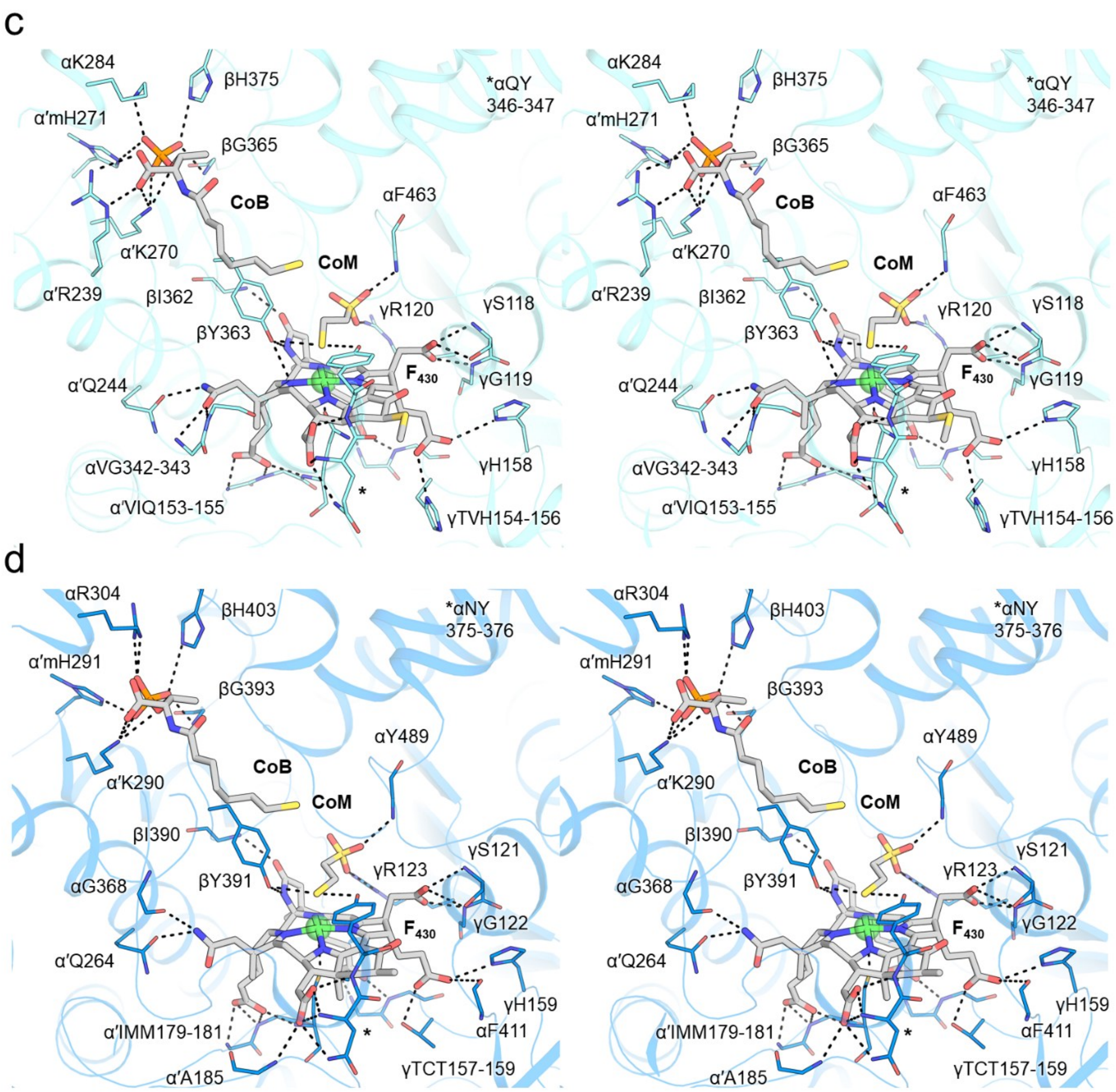


**Supplementary Figure S10. Active site of MCR homologs.** Stereo-view of the MCR active site in (**a**) *M. shengliensis* (7NKG), (**b**) MCR isoform I from *M. marburgensis* (5A0Y), (**c**) ANME-1 (3SQG), and (**d**) '*Ca*. E. thermophilum' (7B1S). The main chains are shown as cartoons, and ligands are shown as sticks coloured according to the atom with red for O, blue for N, orange for P, and yellow for S. Coordinating residues are shown as lines, non-interacting side- or mainchains were omitted. Polar contacts are depicted as black dashes.

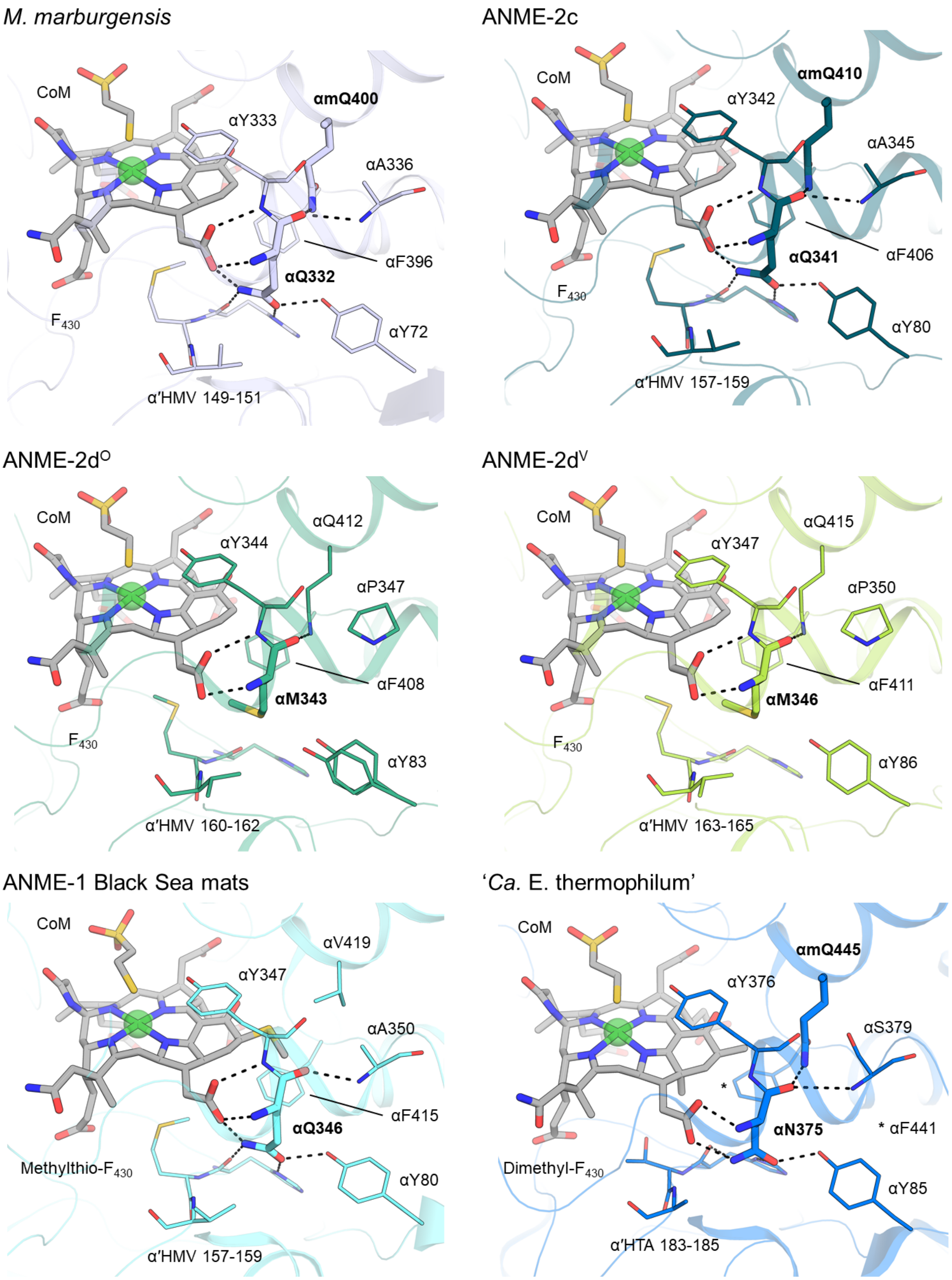


**Supplementary Figure S11. Impact of the glutamine to methionine substitution in the vicinity of the $F_{430}$.** MCR models are shown as a transparent cartoon, and the Gln/Met of interest and the PTM, cofactor $F_{430}$ and CoM are highlighted in sticks. CoB and the heterodisulfide are not shown for clarity. Residues in the close environment of the Gln position 332 and equivalent in homologues are depicted in lines, with polar contacts as grey dashes. Atoms are coloured as follows: red for O, blue for N, orange for P, and yellow for S.

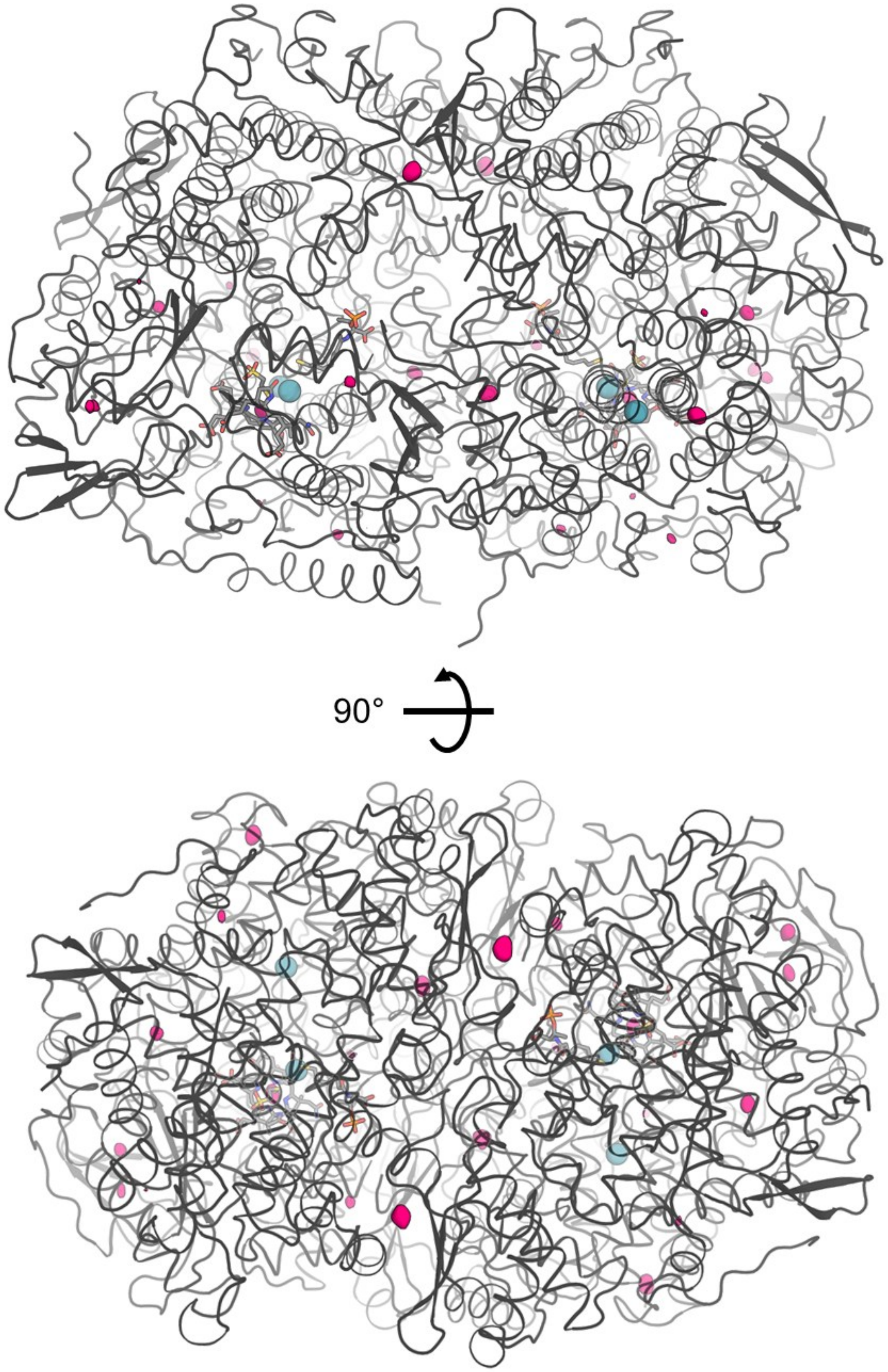


**Supplementary Figure S12. Internal cavities of MCRs.** Anomalous signal of Krypton is depicted by a pink surface contoured to 5 σ. The proteins are shown in cartoons with Xenon positions detected in the gassed structure of '*Ca*. E. thermophilum' (extracted from PDB 7B2C) shown in teal spheres. Coenzymes and $F_{430}$ cofactors are shown as sticks.

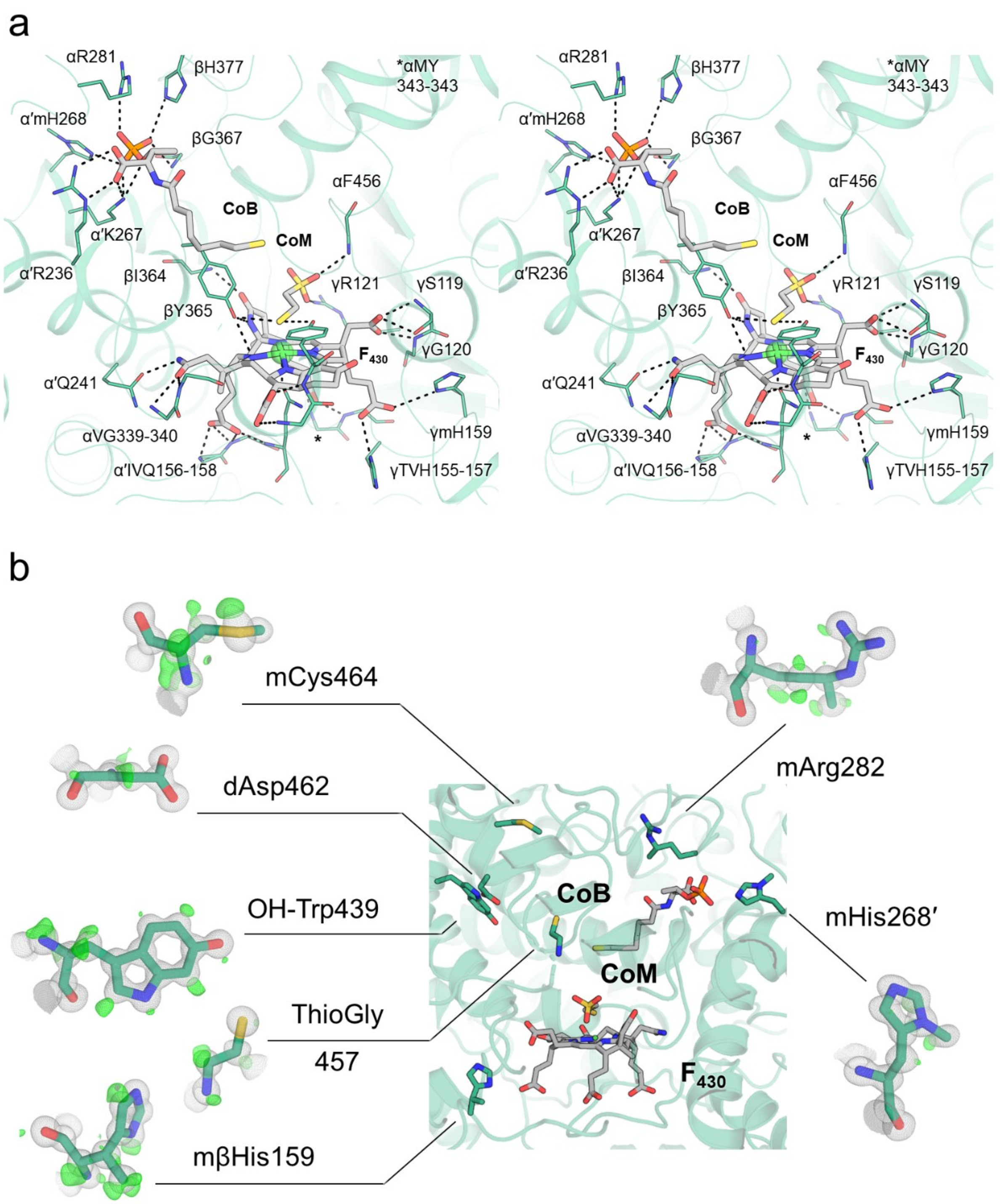


**Supplementary Figure S13. Active site and PTMs in MCR-ANME2d$^{O}$.** (**a**) The active site and (**b**) the PTMs localised in the active site surroundings are depicted in the same way as Figure 3b and Figure 4a, respectively.

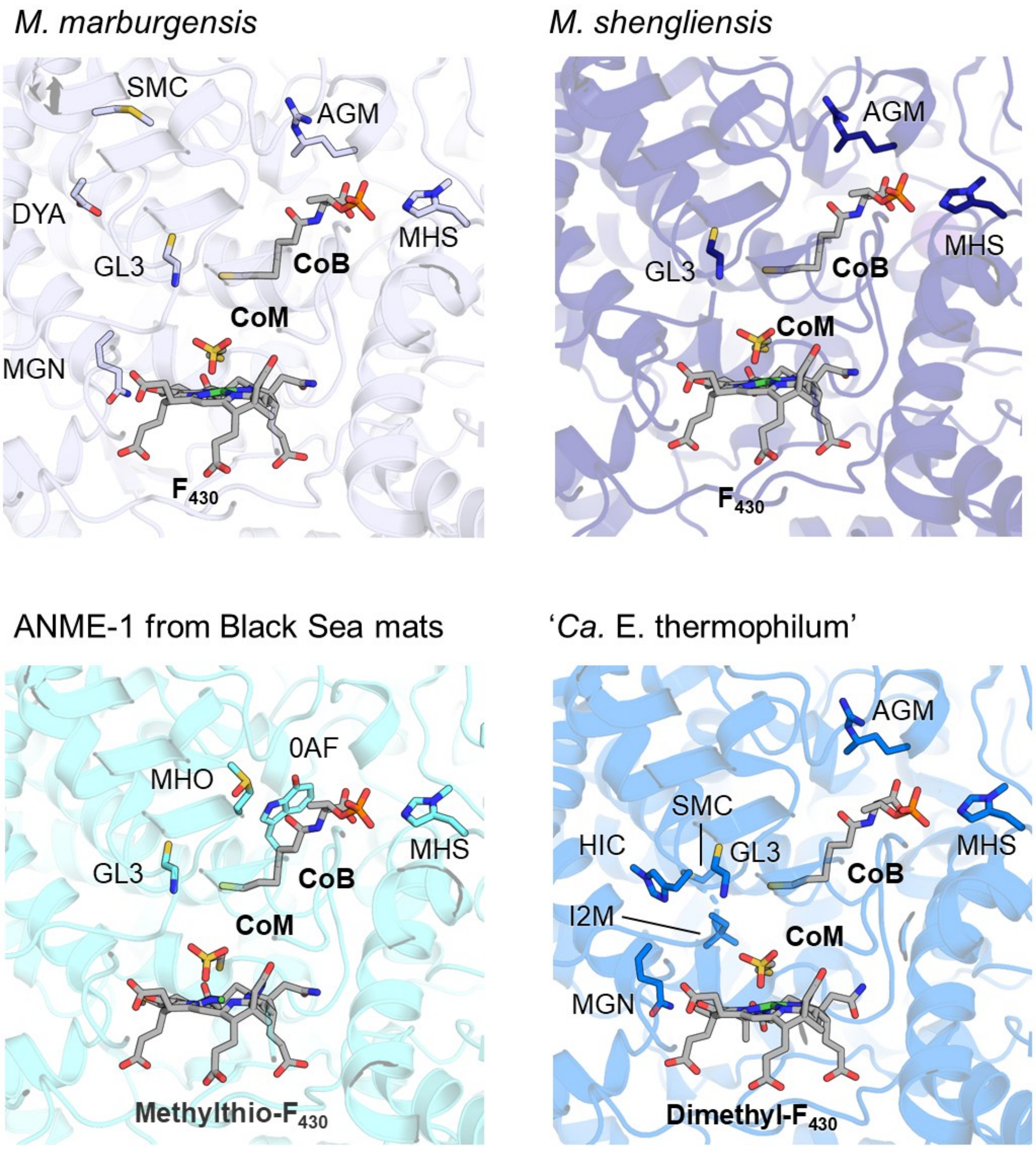


**Supplementary Figure S14. PTMs in MCR homologs.** The main chain is shown as a cartoon, with ligands and PTMs depicted as sticks coloured according to atom (red for O, blue for N, orange for P, yellow for S). Residues are labelled according to the PDB 3-letter ligand codes, with MHS, GL3, AGM, MGN, SMC, DYA, HIC, MHO, and 0AF standing for $N^1$-methylhistidine, thioglycine, 5(*S*)-methylarginine, 2(*S*)-methylglutamine, S-methylcysteine, didehydroaspartate, 4-methylhistidine, S-oxymethionine, and 7-hydroxy-L-tryptophan, respectively.

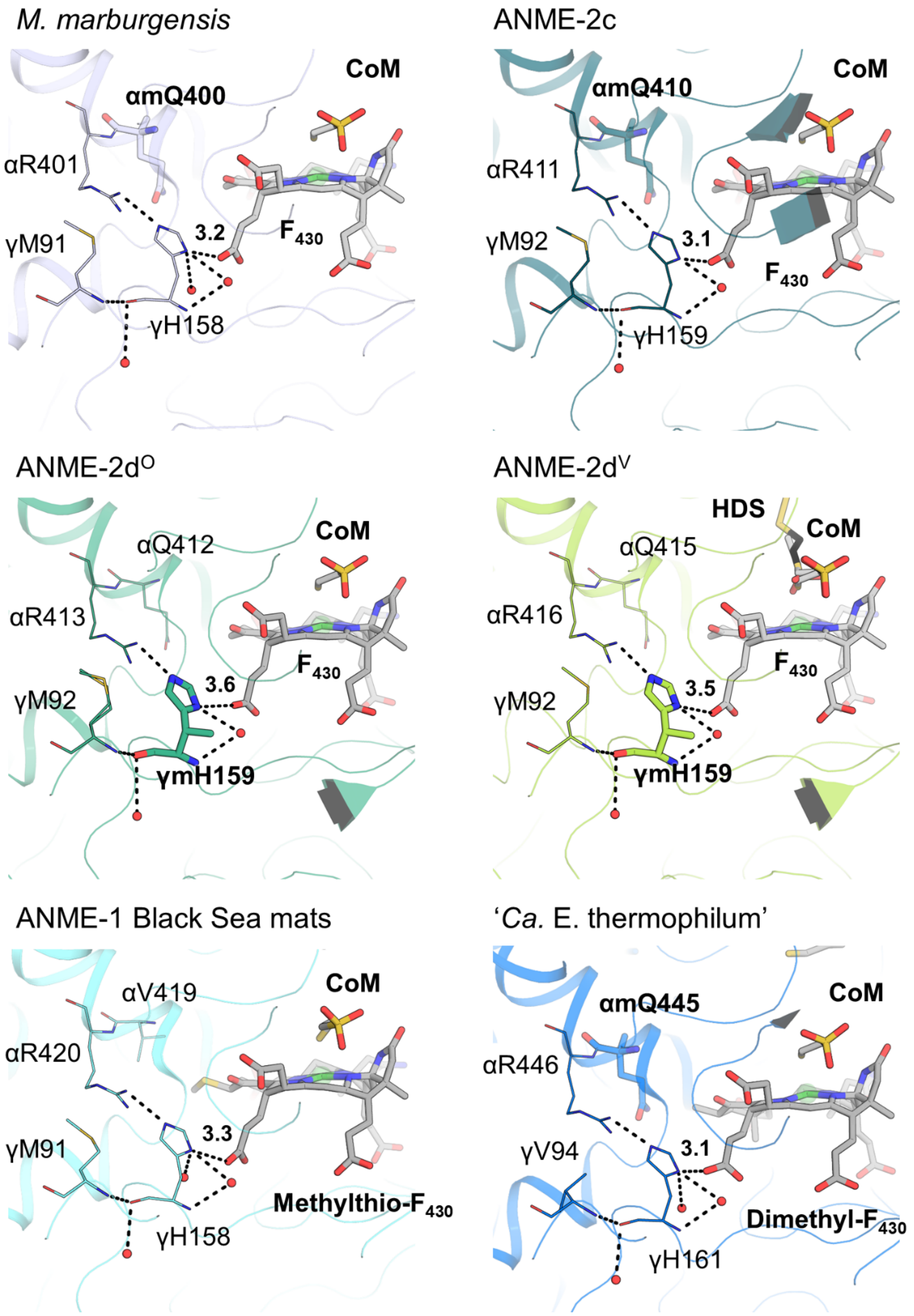


**Supplementary Figure S15. Environment around the 3(*S*)-methylhistidine.** The 3(*S*)-methylhistidine or the equivalent unmodified histidine is shown for the different MCR models. The main chain is depicted as a cartoon, and ligands are shown as sticks coloured as Supplementary Fig. S14. PTMs are shown as sticks and near residues as lines. Polar contacts of histidine/3(*S*)-methylhistidine are depicted as grey dashes. Interacting waters are shown as spheres.

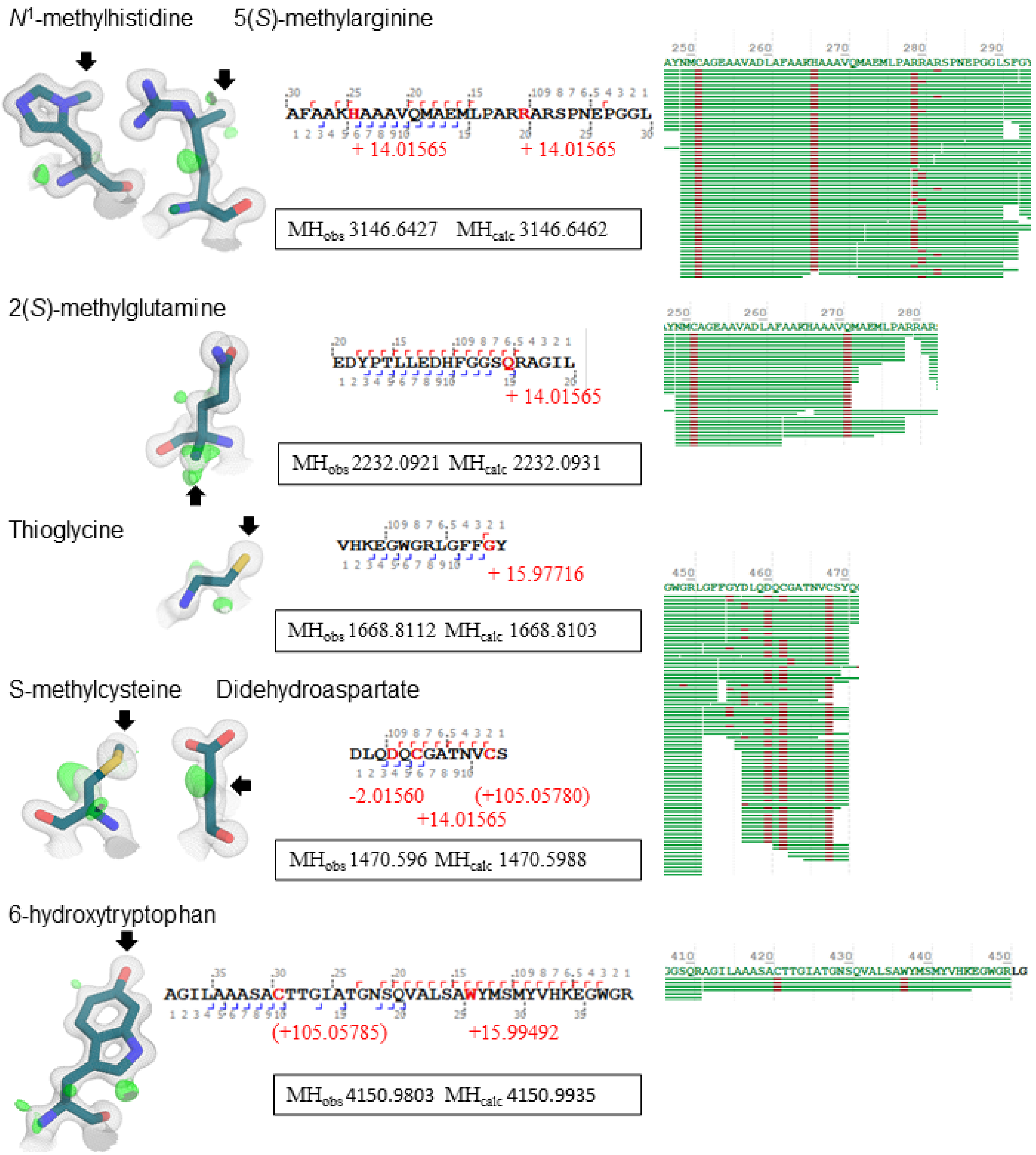


**Supplementary Figure S16. PTMs in ANME-2c MCR detected by LC-MS analysis.** From left to right: Post-translationally modified residue as sticks with the $2F_o-F_c$ (grey mesh) and $F_o-F_c$ map (green mesh) contoured at 1 and 3 σ, respectively. Black arrows highlight the modified position. A representative peptide sequence with fragment ions (hooks) is shown for each PTM. Mass shifts are highlighted in red and shown in brackets if artefactual. The observed ($MH_{obs}$) and calculated ($MH_{calc}$) monoisotopic mass are shown in a box below. An alignment of fragments is shown on the right with modified positions highlighted in red. 6-hydroxytryptophan was treated with chymotrypsin, all others with trypsin.

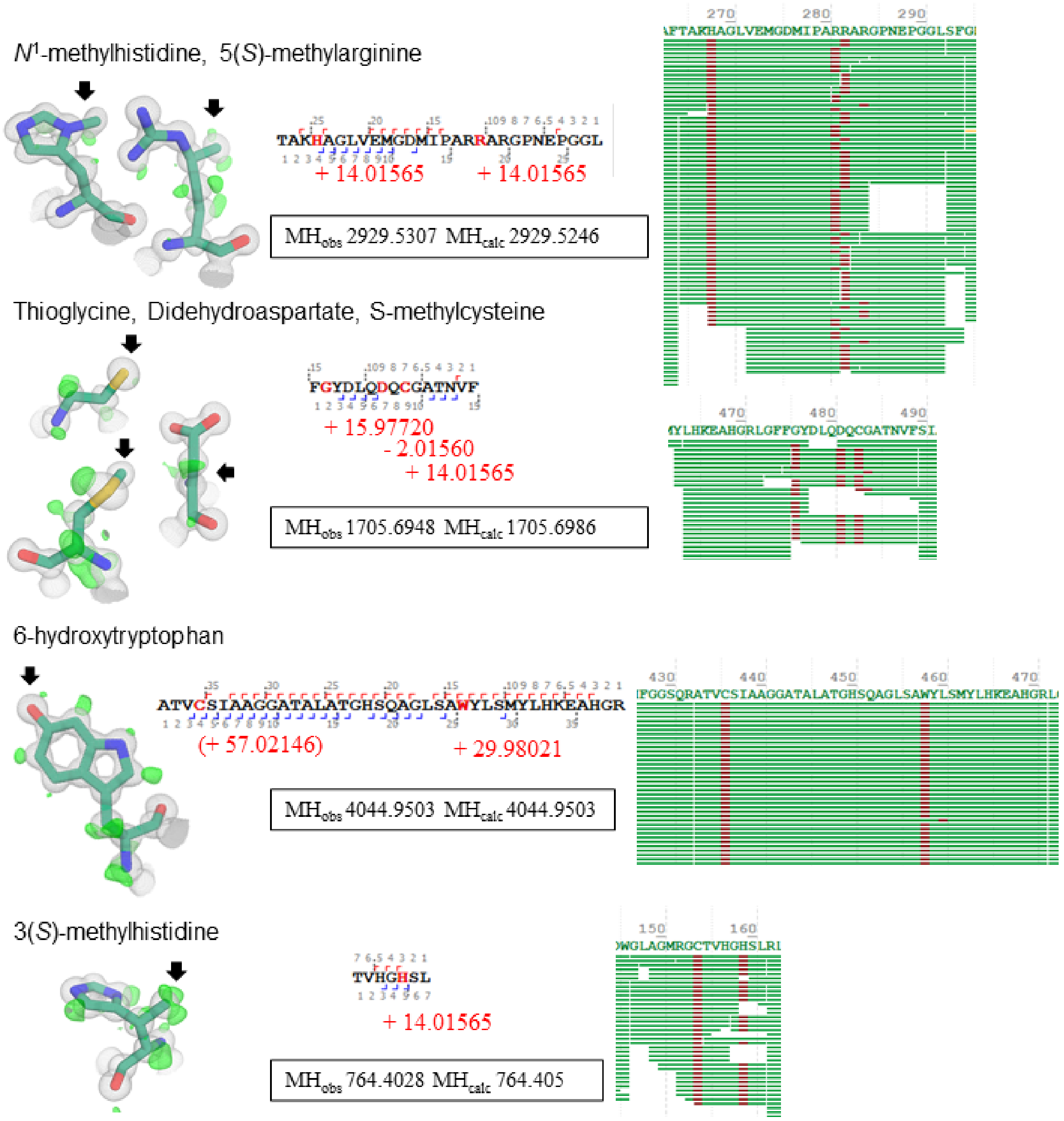


**Supplementary Figure S17. PTMs in ANME-2d$^{O}$ MCR detected by LC-MS analysis.** From left to right: Post-translationally modified residue as sticks with the $2F_o-F_c$ (grey mesh) and $F_o-F_c$ map (green mesh) contoured at 1 and 3 σ, respectively. Black arrows highlight the modified position. A representative peptide sequence with fragment ions (hooks) is shown for each PTM. Mass shifts are highlighted in red and shown in brackets if artefactual. The observed ($MH_{obs}$) and calculated ($MH_{calc}$) monoisotopic mass are shown in a box below. An alignment of fragments is shown on the right with modified positions highlighted in red.

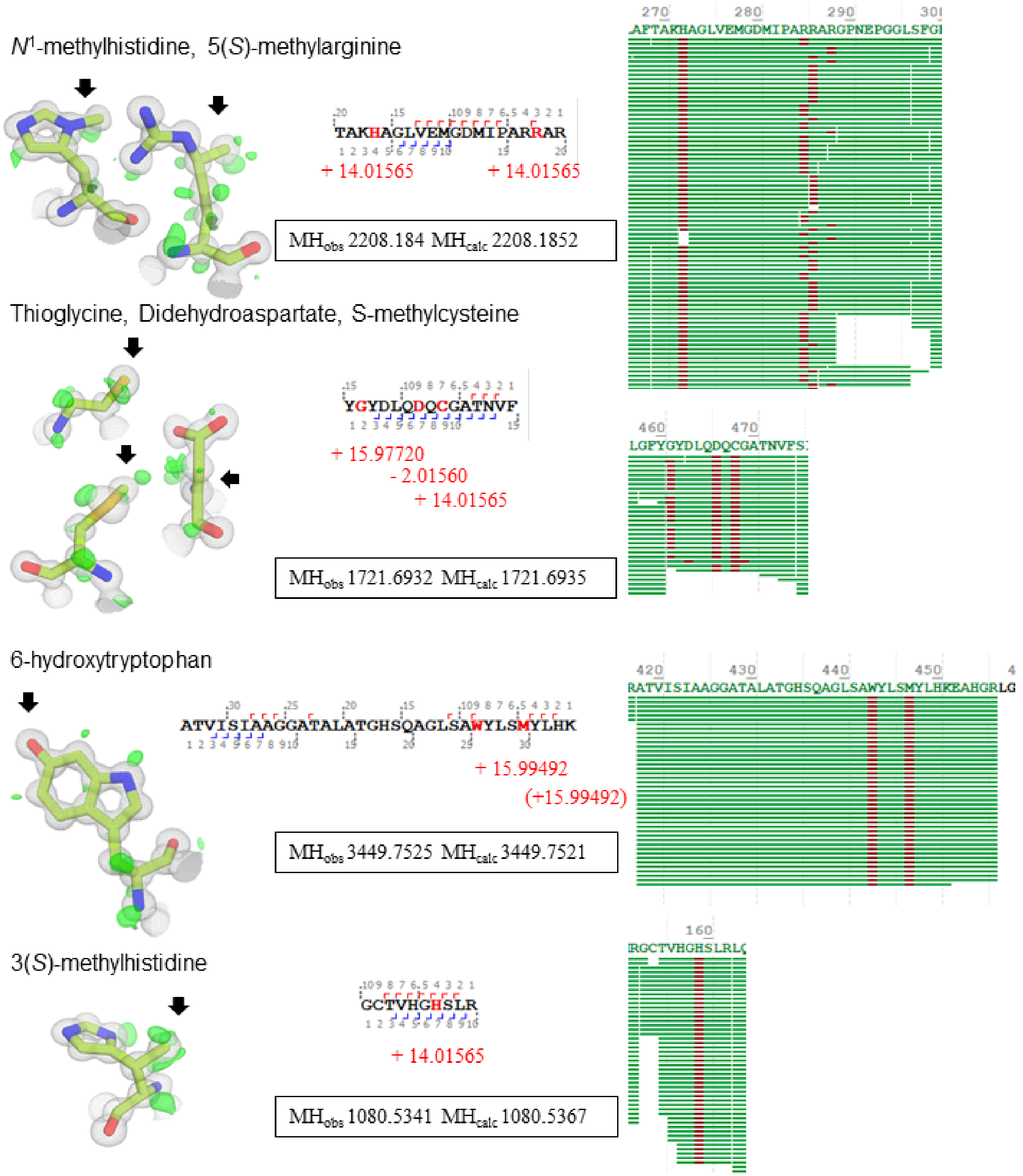


**Supplementary Figure S18. PTMs in ANME-2d$^{V}$ MCR detected by LC-MS analysis.** From left to right: Post-translationally modified residue as sticks with the $2F_o{-}F_c$ (grey mesh) and $F_o{-}F_c$ map (green mesh) contoured at 1 and 3 σ, respectively. Black arrows highlight the modified position. A representative peptide sequence with fragment ions (hooks) is shown for each PTM. Mass shifts are highlighted in red and shown in brackets if artefactual. The observed ($MH_{obs}$) and calculated ($MH_{calc}$) monoisotopic mass are shown in a box below. An alignment of fragments is shown on the right with modified positions highlighted in red.
